# Autoregulation of *RPL7B* by inhibition of a structural splicing enhancer

**DOI:** 10.1101/2025.03.14.643126

**Authors:** David Granas, Ishan Gammadde Hewa, Michael A. White, Gary D. Stormo

**Affiliations:** Department of Genetics and Center for Genome Sciences and Systems Biology Washington University School of Medicine, St Louis, MO

## Abstract

Yeast ribosomal protein gene *RPL7B* is autoregulated by inhibition of splicing. The first intron has a “zipper stem” that brings the 5’ splice site near the branch point and serves as an enhancer of splicing that is required for efficient splicing because it has non-consensus branch point sequence of UGCUAAC. The intron also contains an alternative, and mutually exclusive, structure that is conserved across many yeast species. That conserved structure is a binding site for the Rpl7 protein so that when the protein is in excess over what is required for ribosomes, the protein binds to the conserved structure which eliminates the enhancer structure and represses splicing and gene expression.

## INTRODUCTION

Feedback regulation is common across biology. From product inhibition of enzymatic pathways to autoregulation of gene expression, it is an efficient means of maintaining homeostasis of biological systems [1, 2]. The first discovered example of transcriptional autoregulation was in Phage lambda [3–5]. Two lambda transcription factors (TFs), Repressor and Cro, negatively regulate each other’s expression, but in addition, Repressor initially activates its own expression by binding cooperatively to two adjacent binding sites, and then at higher concentration turns off its own expression by binding to a third site. This positive-negative autoregulation system keeps the concentration of Repressor in a narrow range, poised to become activated at the appropriate signal [6]. Since then, many examples of autoregulation by TFs have been identified [7–9]. Evolution of TF autoregulation is readily accomplished because non-coding DNA evolves rapidly and the occurrence of binding sites for the TF in the appropriate location can be selected if it offers an advantage, which autoregulation often does [10–12].

Post-transcriptional autoregulation has also been observed frequently. In eukaryotic cells, RNA-binding proteins (RBPs) that control gene expression and alternative splicing can evolve to autoregulation, similarly to TFs, by selecting for binding sites for the RBP in appropriate locations in the non-coding regions of the gene [13–15]. Many RBPs function as enhancers or silencers of splicing, often regulating the choice of alternative isoforms of genes, and in many cases regulate their own expression by affecting splicing choices [16]. In bacteria, autoregulation is commonly observed even for proteins that do not normally function as regulators of expression. Most bacterial ribosomal genes are post-transcriptionally regulated either through direct autoregulation, or by being part of an operon that is regulated by one of its constituents [17–20]. Some tRNA synthetase genes are regulated by a mimic of a tRNA structure that occurs near the ribosome binding site, so when the protein is in excess over the amount needed for its normal function, it represses its own expression [21–23]. While ribosomal proteins and tRNA synthetases are both RNA binding proteins, there are other examples of autoregulation of proteins that do not bind RNA as their normal function. First discovered in bacteriophage T4, the ssDNA-binding protein (gene 32) and DNA polymerase (gene 43) are each autoregulated at the translation initiation step, even though neither binds RNA as its normal function [24–29]. Many other examples have been found in bacteria of RNA structures being involved in gene regulation [30, 31]. Because of the enormous potential for RNA to bind to diverse ligands, it can evolve to be an exquisite sensor of cellular states including the concentrations of any functional class of genes. Riboswitches are prime examples of RNAs that have evolved to control the fate of their RNA, either by attenuation, translation or degradation based on the concentration of small molecules in the cell, without any protein requirement [32–37]. The invention of methods such as SELEX further highlights the enormous capability of RNAs to bind to a vast repertoire of ligands, essentially any protein [38–40], which offers the possibility of widespread roles in regulating gene expression [41–43].

Previously, we utilized the yeast GFP-fusion collection [44] to search for yeast ribosomal proteins that showed feedback regulation [45]. For each of those genes we synthesized a copy of the gene encoding the same protein but lacking any potential cis-regulatory elements (“cre-less” versions of the genes), whose expression could be induced from the *GAL1* promoter. Feedback regulation was detected by the ability of the cre-less gene to repress the GFP-tagged wild-type gene. Of the 95 genes tested, 30 showed at least a 3-fold reduction in GFP expression after induction of the cre-less gene. We delved into the mechanism of regulation of *RPL1B*, a gene that was repressed 10-fold upon induction. We found that it contains an RNA sequence and structure immediately 5’ of the initiation codon that is conserved from bacteria to archaea to eukaryotes in both the binding sites of the protein on rRNA and in the mRNA of the gene [46–49]. We also showed that disrupting that conserved sequence or structure eliminated regulation. Those results show that *RPL1B* autoregulation in yeast is very similar to many examples of bacterial autoregulation, where binding of the protein inhibits translation initiation.

In addition to regulation by simply blocking translation initiation, yeast can autoregulate by inhibition of splicing. Several examples of autoregulation of ribosomal protein expression by inhibition of splicing are known: *RPL22B, RPS41B, RPL30* and *RPS9A* and *B*, are all regulated in that manner [50–55]. The non-ribosomal protein Yra1, which is involved in nuclear export, is also autoregulated by inhibition of splicing [56]. Most yeast genes do not contain introns, but ribosomal protein genes are the exception, with 76% of them containing introns. Only nine genes in yeast contain two introns, three of which are ribosomal protein genes [57]. The length distribution of introns in yeast is bimodal, with ribosomal genes having mean intron lengths of about 400 bases, whereas non-ribosomal introns are generally much shorter, with an average length closer to 100 bases [58]. Each of the yeast introns has been deleted, and many of those strains have growth phenotypes under some conditions [57, 59–62]. The secondary structure of yeast pre-mRNAs has long been recognized as important in influencing splicing efficiency [63–67]. A recent study of yeast intronic RNA structures, using a combination of *in vivo* structure probing experiments and computational prediction, showed that most introns are highly structured, and it is common for there to be “zipper stems” that effectively shorten the intron length and bring the 5’ splice site close to the branch point (BP) [68]. In *RPS17B* it was demonstrated that such an RNA structure served as an enhancer for splicing [69]. Computational analyses have suggested that the intron length between the 5’ SS and BP, taking into account secondary structures that effectively shorten that distance, is correlated with splicing efficiency with longer introns being less efficiently spliced [67]

In this paper we focus on *RPL7B* (also known as *uL30* [70]) because our previous work showed it is likely to be autoregulated [45] and because the *RPL7B* gene has several interesting features that we suspected could be involved in the regulation. *RPL7B* and *RPL7A* contain two introns [71], which is very rare in yeast, occurring in only one other ribosomal protein gene [57, 58]. Both paralogs contain a snoRNA in the second intron. The first intron of both paralogs contains a predicted RNA secondary structure that is conserved across many yeast species [72, 73] and the first intron also contains the unusual branch point (BP) sequence UGCUAAC, which only occurs in one other yeast gene [58]. The deviation from the consensus BP sequence of UACUAAC has been shown in other work to cause inefficient splicing [74, 75].

## MATERIALS and METHODS

### Yeast strains

Plasmids MJB1-*RPL7A* and MJB1-*RPL7B* were created to express the cre-less versions of these genes as proteins fused to mCherry at the C-terminus, and under control of the inducible GAL1 promoter. “Cre-less” refers to the fact that the 5’ and 3’ UTRs have been replaced, the introns have been removed, and the codons have been shuffled to give the same protein sequence from a different mRNA sequence. These cre-less sequences were synthesized by Synbio and cloned into the MJB1 vector (previously described in Roy et al., 2020) using NEB’s HiFi DNA Assembly. These plasmids contain the URA3 gene for selection in ura-yeast strains.

The Rpl7a-zz protein was expressed from *the RPL7A* plasmid from the movable ORF collection [76]. This plasmid expresses Rpl7a with 3 C-terminal tags: 6x-His, HA, and ZZ Protein A. It also contains URA3 for selection in ura-yeast strains.

Yeast strains were constructed to express the eGFP gene containing different intron variants. The wild-type introns were cloned into the plasmid BAC690-TEF (previously described in Roy et al., 2020) using NEB’s HiFi DNA Assembly. Intron variants were made using NEB’s Site-Directed Mutagenesis Kit. A PCR amplicon was then generated from the plasmid containing the TEF2 promoter, eGFP with intron(s), ADH1 terminator, kanamycin/G418 resistance marker, and homology arms for integration into the yeast genome at the location of the dubious ORF YBR032W.

Yeast transformation was done using the LiAc/SS carrier DNA/PEG method. Transformants were selected on YPD plates with 400 μg/mL G418 and verified by sequencing.

### GFP assays

Liquid cultures were inoculated and grown overnight in 400 µL SC-URA with 2% raffinose in 96-deep-well plates at 30°C. Overnight cultures were diluted into both SC-URA with 2% raffinose (uninduced) and SD-URA with 2% raffinose and 0.2% galactose (induced). Cells were grown at 30°C for 10 hours. 200 µL cultures were transferred to 96-well plates and assayed on a CytoFLEX S (Beckman Coulter). The live cells were gated and 10,000 events were acquired.

### Assaying Rpl7-zz protein binding to the intron sequences

The movable ORF *RPL7A* plasmid was transformed into the strains containing the wild-type and S1 versions of *RPL7B* intron upstream of GFP. Strains were inoculated separately into 3 mL of SC-URA medium supplemented with 2% glucose and incubated overnight at 30°C. A 100 µL aliquot from each overnight culture was transferred into 50 mL of pre-warmed SC-URA medium containing 2% raffinose and incubated at 30°C for 6 hours. Cultures were subsequently diluted and transferred into 200 mL of pre-warmed SC-URA medium supplemented with 2% raffinose to an initial OD_600_ of 0.02 and they were grown at 30°C until OD_600_ 0.7–0.8. From each culture, 100 mL was transferred into sterile 250 mL conical centrifuge bottles, placed on ice, and labelled as time 0 (uninduced). The remaining 100 mL of each culture was combined with 50 mL of pre-warmed SC-URA medium containing 6% galactose and incubated for 15 min to induce Rpl7a-zz expression. A volume of 150 mL aliquot of each galactose-induced culture was transferred to sterile 250 mL conical centrifuge bottles, placed on ice, and labelled as time 15 (induced). Samples were subjected to 1,200 mJ/cm^2^ of 254 nm UV irradiation using a Stratalinker UV Crosslinker 2400 as described in [77]. Cells were pelleted by centrifugation at 1,500 g, 4°C, for 5 min, and resuspended in 200 µL of lysis buffer (20 mM HEPES, pH 7.4, 2 mM Mg(OAc)_2_, 100 mM KOAc, 100 µg/mL cycloheximide, 0.5 mM DTT, and 1× protease inhibitor cocktail [Sigma]). An equal volume of glass beads (500 µm, BioSpec) was added, and cells were lysed by vortexing at 4°C for 15 min. Lysates were centrifuged at 10,000 rpm, 4°C, for 5 min, and the supernatants were collected in new 1.5 mL centrifuge tubes. A 200 µL aliquot of lysate from each sample was mixed with an equal volume of 2× ice-cold binding buffer (100 mM Tris-HCl, pH 7.5, 24 mM Mg(OAc)_2_, 1 mM DTT, 1 mM PMSF, 1 U/mL RNase inhibitor [NEB]) and incubated with 700 µL of IgG Sepharose (Cytiva) in 1.5 mL microcentrifuge tubes. Binding was carried out at 4°C for 1 h with constant gentle rocking. Resins were washed three times with 800 µL of ice-cold 1× wash buffer (50 mM Tris-HCl, pH 7.5, 100 mM KCl, 12 mM Mg(OAc)_2_, 1 mM DTT, 1 mM PMSF), followed by DNase I treatment in 500 µL of nuclease-free water containing 10 U of DNase I (NEB) in 1× DNase I reaction buffer at 37°C for 10 min. Resins were then washed three additional times with 800 µL of ice-cold 1× wash buffer, and the final pellet was resuspended in 500 µL TRIzol (ThermoFisher) for RNA extraction according to the manufacturer’s protocol. Total RNA was extracted from a separate isolate of the T0 samples by the addition of 500 µl TRIzol. RNA was then purified using the Monarch Spin RNA Cleanup Kit (10 µg) (NEB) and eluted in 10 µL of nuclease-free water and quantified using a NanoPhotometer® NP50. For cDNA synthesis, SuperScript™ IV First-Strand Synthesis System (Invitrogen) was used with the following gene-specific primers: RIP-GFP_cDNA for GFP, RPL7A_cDNA for RPL7A, and RPL7B_pull_cDNA for RPL7B (Supplemental Table S1). GFP cDNA from total RNA (T), wild-type, and S1 was PCR-amplified using T7_hooks_structure_F and T7_hooks_structure_R primers. Similarly, *RPL7A* cDNA was amplified using *RPL7A*_Intron_F and *RPL7A*_Intron_R, while *RPL7B* cDNA was amplified using T7_hooks_structure_F and T7_hooks_structure_R primers using Q5 Hot Start High-Fidelity DNA Polymerase (NEB). The final PCR products were analyzed by electrophoresis on a 3% agarose gel.

## RESULTS

To test if the introns of *RPLA* and *RPL7B* are sufficient as the cis-regulatory elements (CREs) for regulation, we inserted them into a chromosomal copy of GFP under control of the constitutive *TEF2* promoter [78]. We drove expression of the cre-less Rpl7a protein from a plasmid under the control of the *GAL1* promoter [45]. We compared the GFP expression levels by flow cytometry with and without galactose induction of the Rpl7a protein. Figure 1a is a schematic of the experimental design where “Insert” refers to the intron variants that were tested for *cis*-regulatory activity. Figure 1b shows the fluorescence levels of cells without *GFP*, as background, and Figures 1 c-e for cells in which both introns or only intron 1 or intron 2 of *RPL7B* are inserted into the *GFP* gene, with and without induction. Regulation of about 7-fold (0.85 change in log(fluorescence)) is observed when *GFP* contains both introns and only intron 1. With only intron 2 there is only a slight reduction in expression after induction. Figure 1 f-g shows that there is no significant regulation when both introns or only intron 1 of *RPL7A* are present. We therefore focus on testing variants of intron 1 of *RPL7B* to elucidate the mechanism of regulation. The paralogous proteins Rpl7a and Rpl7b differ at five amino acids and endow ribosomes with somewhat different properties [79–83] so we tested the Rpl7b protein for regulatory function by expressing a cre-less version of the gene. We found it to be equivalent to Rpl7a protein in that it regulates only *RPL7B* and not *RPL7A* (Supplemental Fig. S1), so all further work was performed with the Rpl7a protein. (Expression values for all tested variants are shown in Supplemental Table S2.)

**Figure 1.**
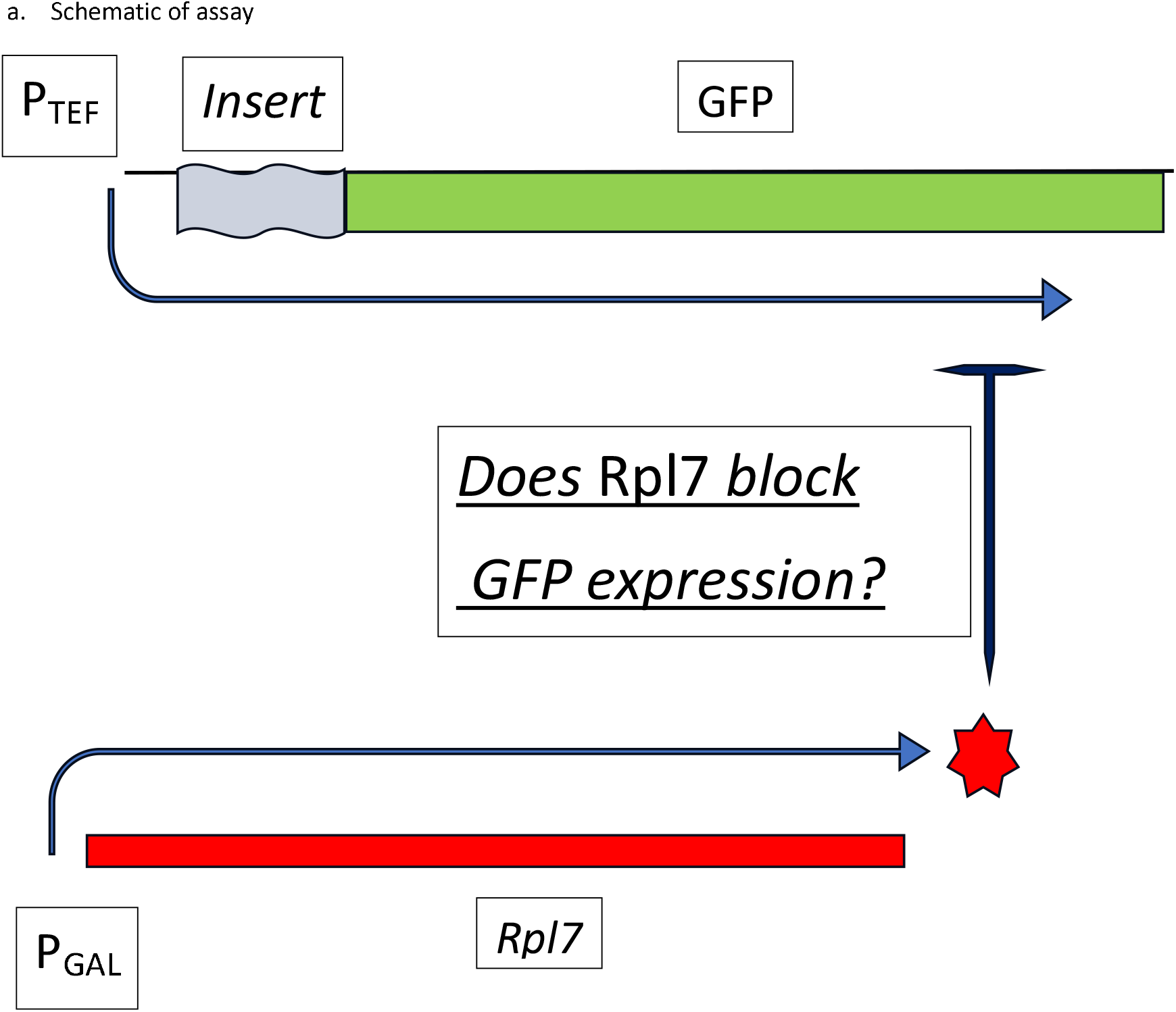
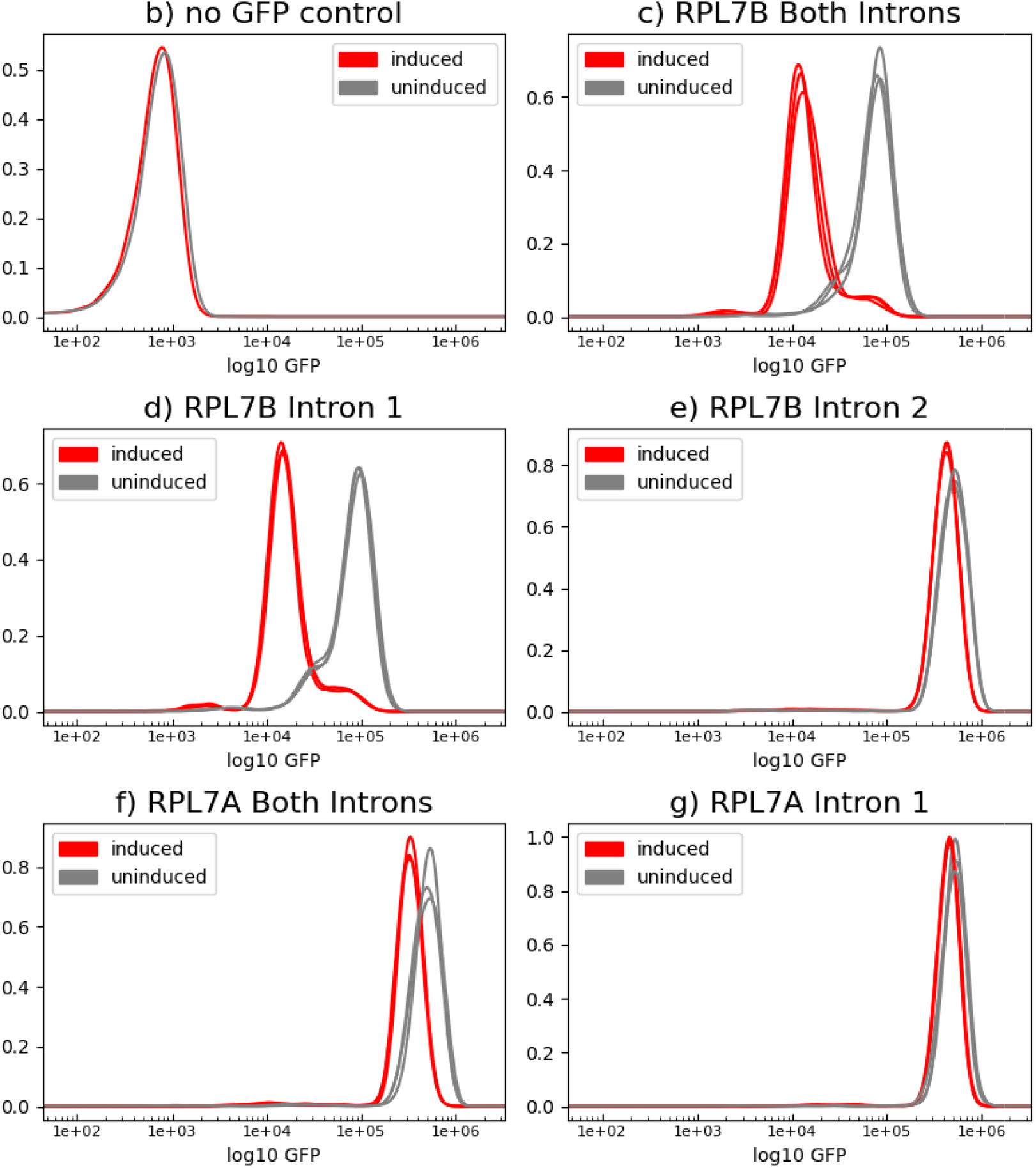
Experimental Design and Results a. Schematic of experimental design. Rpl7 protein is induced by adding galactose to the medium. The *Rpl7a* gene is the *cre*-less version that has eliminated any potential *cis-regulatory elements* [45]. “insert” refers to different potential regulatory elements inserted into the GFP gene. Proper splicing of those inserted elements leads to GFP protein production. b.-f. Flow-cytometry histograms of fluorescence signals at 525 nm wavelength to detect GFP protein levels. Replicate experiments are overlaid and can be seen to be nearly identical. c. yeast cells in the absence of GFP gene to show background fluorescence. d. The insert contains both introns 1 and 2, and the short exon between them, of *Rpl7b*, under conditions of both Rpl7 induction and uninduced. e. The insert contains only intron 1 of *Rpl7b*. f. The insert contains only intron 2 of *Rpl7b*. g. The insert contains introns 1 and 2 of *Rpl7a*. e. The insert contains only intron 1 of Rpl7a.

We sought to identify the regions of intron 1 that are necessary for regulation by testing whether deletions eliminated regulation. Figure 2a shows a schematic of *RPL7B* intron 1 with various deletions indicated. The entire intron is 409 bases long with the first U of the UGCUAAC BP at position 382 after the 5’ splice site. There is a predicted secondary structure that is conserved between 36 species of fungi from bases 200 to 355 within the intron [73], which we refer to as the Hooks structure (Figure 2b). U1-U3 are about 63-base segments upstream of the Hooks structure, but after the 5’ splice site sequence, that were independently deleted. D1 is the segment between the Hooks structure and the BP that was deleted. S1-S4 are various deletions of the Hooks structure: S1 deletes the entire structure and S2-S4 delete independent stems or loops, as shown in Figure 2b. We also replaced the wild-type BP sequence of UGCUAAC with the consensus UACUAAC in each of the deletion mutants to assess its role in regulation.

**Figure 2.**
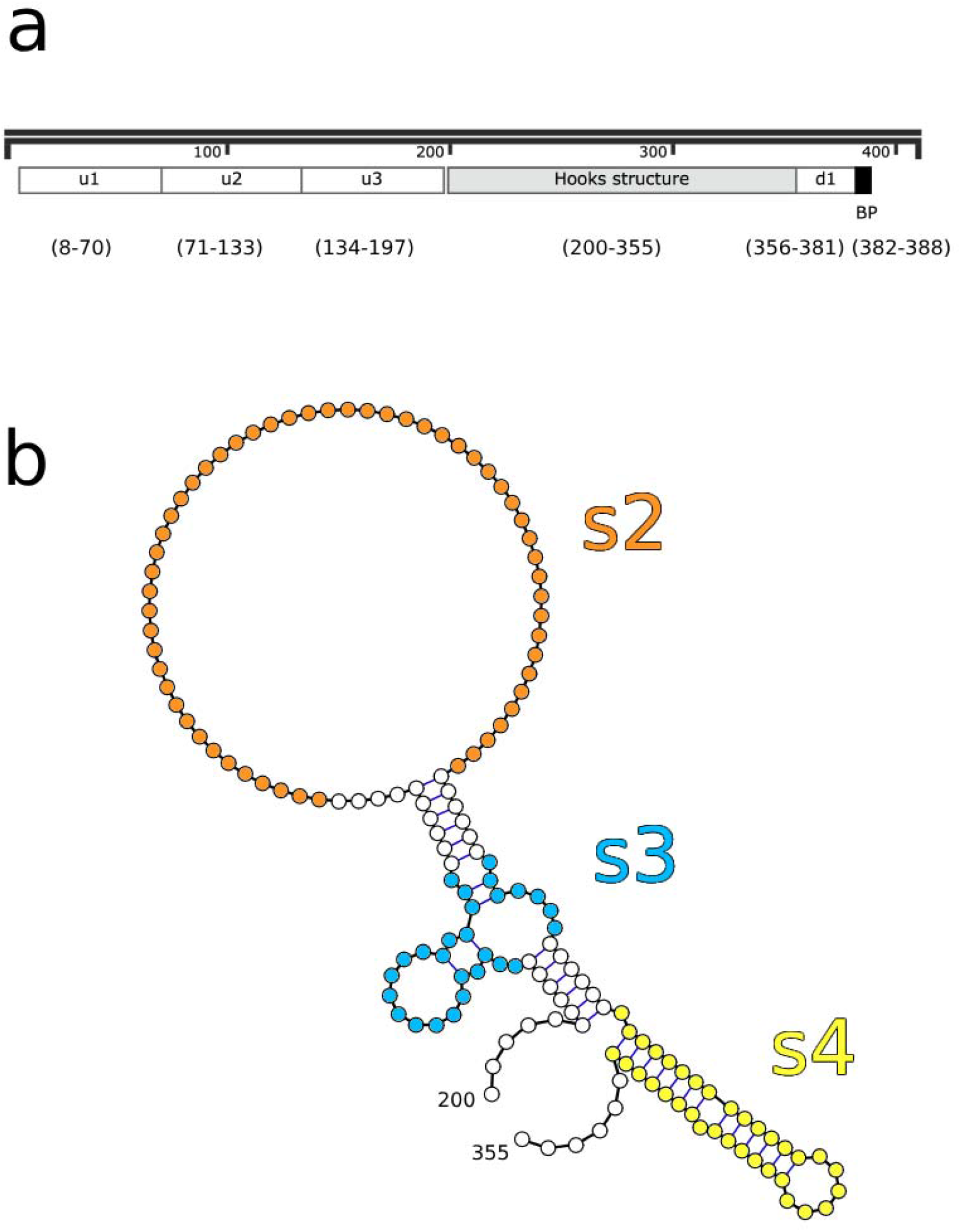
Schematic of intron 1 with deletions. a. The 5’ splice site is at position 1 of the intron and the 3’ splice site is at position 409. The BP (BP) sequence UGCUAAC is at positions 382-388. The positions of deletions u1-u3 and d1 are indicated, as is the position of the conserved Hooks structure. b. A diagram of the conserved Hooks structure with the deletions s2-s4 indicated in colors.

Figure 3 summarizes the primary results. The names of each mutant are listed across the top of the figure. The vertical axis is the log of the fluorescence measurement with 0 defined as the wild-type intron 1 in the uninduced (not repressed) state. Each the four vertices of each quadrilateral represent the induction and BP states. The upper left (star in the wild-type panel) indicates the GFP expression with the wild-type BP without induction of the cre-less Rpl7a protein. The lower left (triangle in the wild-type panel) is expression with the wild-type BP after induction. In every case induction of Rpl7a reduces GFP expression, but the amount varies widely. The upper right vertex of the quadrilateral (square in wild-type panel) is the fluorescence when the wild-type BP sequence is replaced by the consensus sequence, in the absence of repression. In every case the consensus BP leads to higher expression, but the amount of change varies widely. The lower right vertex (circle in wild-type panel) is the fluorescence after induction of introns containing the consensus BP sequence.

**Figure 3.**
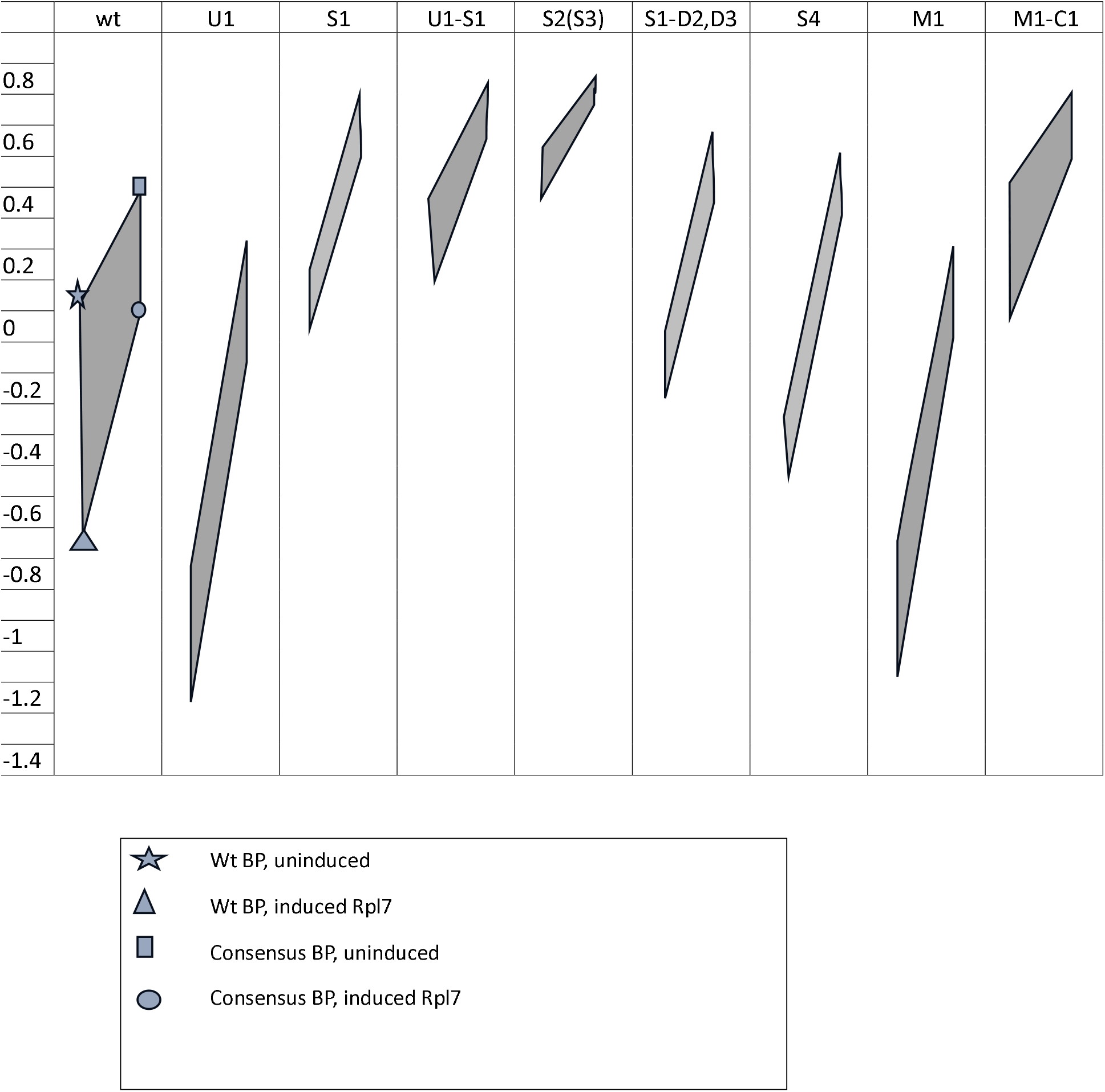
Plots of fluorescence values for different intron inserts. The vertical lines show the change upon induction of the Rpl7 protein, and the horizontal lines in each quadrilateral show the changes upon substitution of the consensus BP (UACUAAC) for the wild-type BP (UGCUAAC).

Deletion U1 greatly reduces expression in the uninduced state, even lower than the repressed wild-type intron. It is still regulated, to a reduced extent. Replacing the wild-type BP sequence with the consensus restores expression, even slightly above the wild-type, and repression is maintained at the reduced level. U2 and U3 are not shown. They both exhibited about 2-fold higher expression than wild-type, and regulation was nearly the same as wild-type. We suspect these mutants have increased expression simply due to the shorter intron, and they were not studied further. The fact that the sequence deleted in U1 is necessary for high expression only with the wild-type BP suggests it contains a splice enhancer element that compensates for the weak BP.

To test the importance of the Hooks structure for regulation, the entire structure was deleted in S1. It still contains the splice enhancer region so is expressed at slightly higher level than wild-type, but it has lost most of the regulation. With that deletion, expression is repressed about 2-fold upon induction of Rpl7, instead of the 7-fold for wild-type. The consensus BP increases expression even more than in wild-type, perhaps due to the shortening of the intron. The combination of deletions U1 and S1 is similar to S1 alone, but somewhat higher expression even in the absence of the enhancer region, possibly because it is much shorter than the other introns tested. Deletions S2 and S3 remove the first and second stem-loops of the Hooks structure, and have similar effects to S1, with increased expression but reduced regulation. Deletion S4, which removes the third stem of the Hooks structure, has a much more severe effect than any of the other S-deletions, even S1 that removes the entire Hooks structure. It is also nearly insensitive to repression and is rescued to a larger degree upon replacement of the BP with a consensus sequence.

Deletion D1 removes the entire sequence between the Hooks structure and the BP. It has essentially no expression, only 0.1 above the background level, and is insensitive to either repression or replacement of the BP (not shown). Although D1 retains the entire canonical BP sequence (UGCUAAC and UACUAAC) it must remove some critical element for splicing. We made smaller deletions, D2-D4 each with nine bases deleted, in conjunction with S1, to determine if they are required for the enhancer activity. S1-D4 is essentially the same as S1 alone (not shown), but S1-D2 and S1-D3 each reduce expression about 0.3 below S1 alone, and show a larger increase in expression with the consensus BP. A possible explanation for these results is that the sequences under U1 and D2-D3 form a secondary structure that serves as a splicing enhancer which is required for high level expression with the wild-type BP.

### Enhancer and regulatory structures

In a study of the *RPL7A* intron, Howe and Ares [84] proposed a secondary structure between the 5’ end of the intron and the region just 5’ of the BP that effectively shortens the distances between the 5’ splice site and the BP. More recent work [68], using *in vivo* chemical probing and structure prediction, confirmed that structure and extended it to a more stable structure. The sequences covered by U1 and D2-D3 in *RPL7B* differ substantially from the sequences in *RPL7A*, but a similar possible structure can be formed in the *RPL7B* intron. Figure 4a outlines our model for a secondary structure that brings the 5’ splice near the BP and can serve as a splice enhancer. Figure 4b shows that structure in detail on the left. The bases in yellow, in Figures 4a and b, are the same bases that can form alternative structures. The structure in the lower part of Figure 4a, and shown in detail on the right in Figure 4b, is part of the conserved Hooks structure. We postulate that the Rpl7 protein binds to the Hooks structure, shifting the equilibrium towards that structure and away from the enhancer structure, thereby reducing expression. The structure on the right is probably just a portion of the binding site for Rpl7 protein, as deletion of the other stems, S2 and S3, also greatly reduced the level of repression. The reason that deletion S4 reduces expression even more than S1, is that while they both remove the 3’ portion of the enhancer structure, S1 also reduces the size of the intron by over 150 bases, whereas S4 only removes 40 bases, leaving the intron still quite long compared to most yeast introns. In both cases, and especially in S4, changing the BP to a consensus sequence eliminates the need for the enhancer structure, giving rise to higher expression than the wild-type intron.

**Figure 4.**
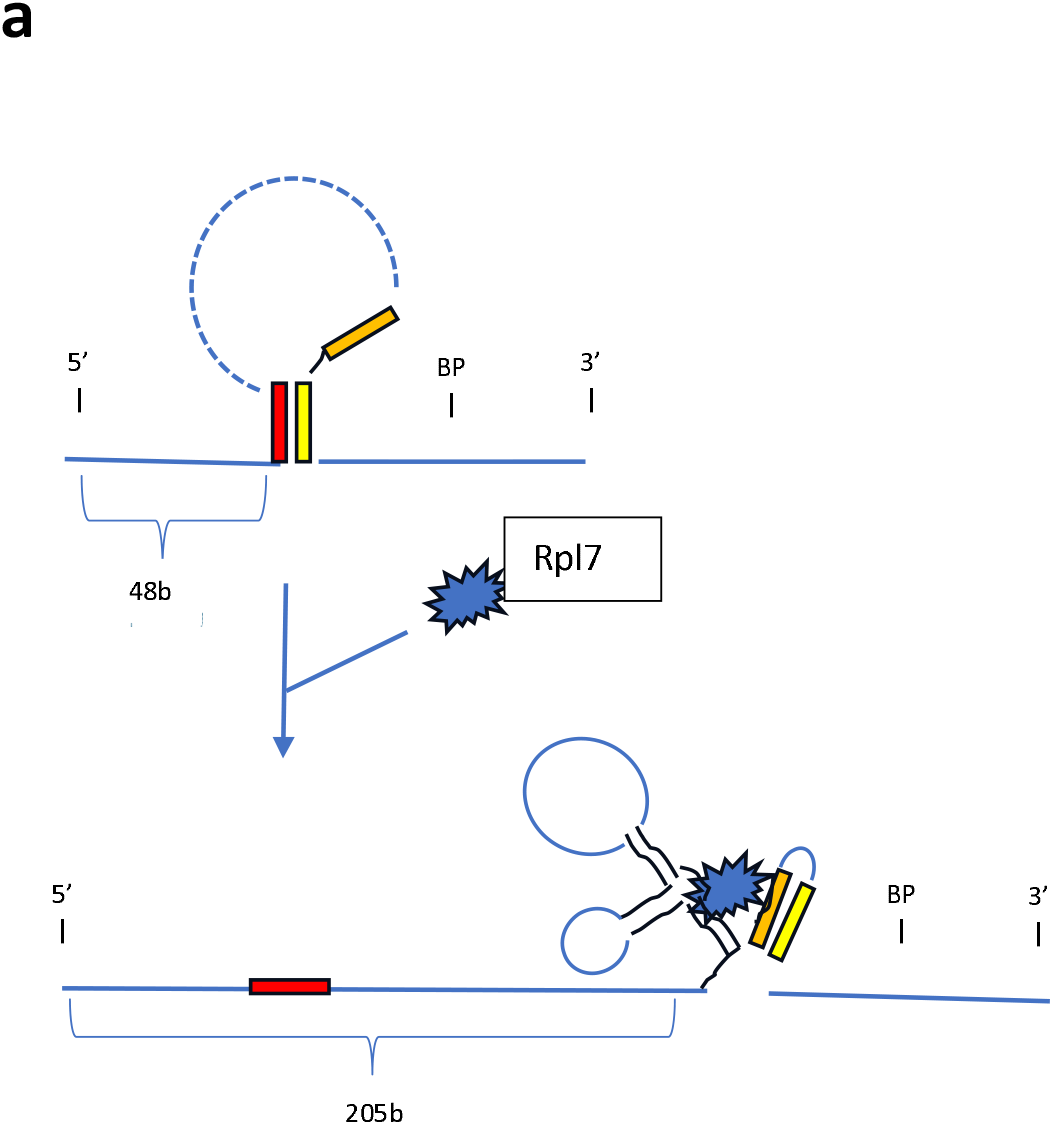
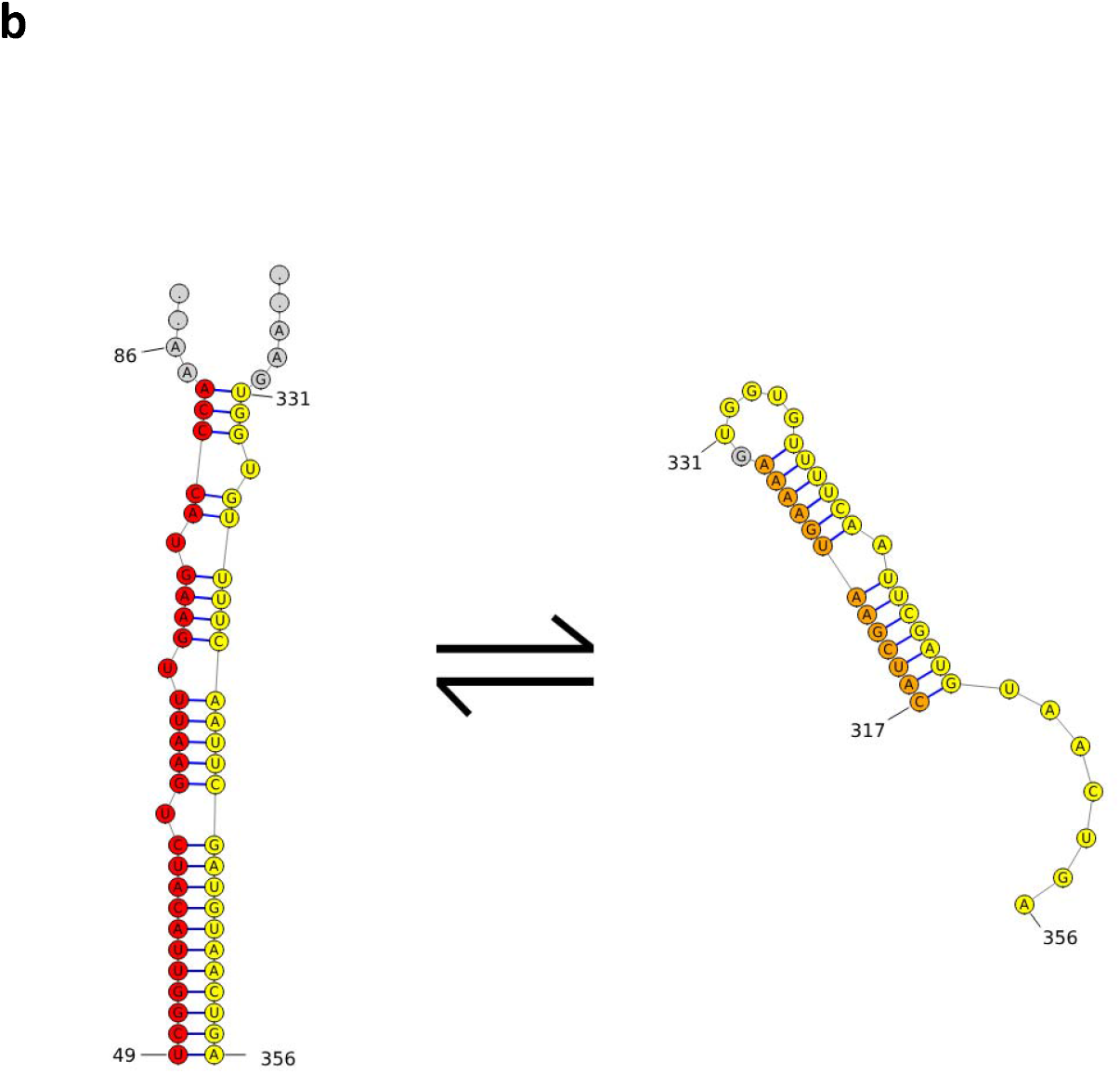
The model for splicing regulation. a. The intron can form a structure between sequences at near the 5’ end (red) and sequences near the BP (yellow) that serves as a splicing enhancer. Rpl7 protein can bind to an alternative structure (Hooks structure) which include the orange and yellow sequences. This eliminates the enhancer structure and makes the BP more distant from the 5’ end of the intron. b. The figure on the left is the proposed structure of the *RPL7B* splicing enhancer, that pairs positions 49-84 with positions 331-356 of intron 1. The structure on the right is stem 3 of the Hooks structure (deleted in s4, shown in Figure 2b). The bases shown in yellow are the same bases shown in mutually exclusive, alternative structures.

To further test this model, instead of deleting the large segment U1, we changed just the first ten bases shown in the left of Figure 4b, from positions 49 to 58, to a non-complementary sequence. This mutant would reduce the stability of the enhancer structure and shift the equilibrium to the structure on the right. This intron mutation is labeled M1 in Figure 3, which shows that it behaves nearly identically to the large deletion U1. We then altered the sequence at the 3’ end of yellow sequence, to compensate for the M1 mutation and reform the enhancer structure, while leaving the right structure unchanged. This version is labeled M1-C1 in Figure 3, which shows that it restores behavior of the intron to near wild-type activity. In fact, it expresses GFP at a somewhat higher level than the wild-type intron, and shows somewhat reduced repression, probably because the mutated and compensated structure is predicted to be somewhat more stable than the wild-type enhancer structure.

### Direct protein binding to the intron

The model requires that the Rpl7 protein bind to the intron to inhibit splicing. To test this, we overexpressed a version of Rpl7a protein fused to a protein A ZZ domain tag (Rpl7a-zz) [76] in the strains containing either the wild-type intron or the Hooks deletion (S1) in *GFP*. Expression of Rpl7a-zz was induced with galactose and samples were taken at 0 and 15 minutes after induction. After UV crosslinking, Rpl7a-zz was isolated by immunoprecipitation with IgG resin, followed by reverse transcription of the bound RNA. We then used gene-specific PCR primers to determine whether the *GFP* or native *RPL7A* or *B* introns were bound by Rpl7a-zz (Figure 5). No PCR product was present at time 0, as expected because the tagged protein has not yet been expressed. At 15 minutes, the wild-type intron of *GFP* is bound by the protein, but the S1 version of the intron, with the Hooks structure deleted, is not bound. The *RPL7A* intron is not bound in either strain, consistent with that gene not being regulated. The *RPL7B* intron is bound in both wild-type and S1 strains, which is consistent with the model. The lanes labeled T are for total RNA extracted from the respective strains and show that all the intron sequences are present and amplified specifically by the designed primers.

**Figure 5.**
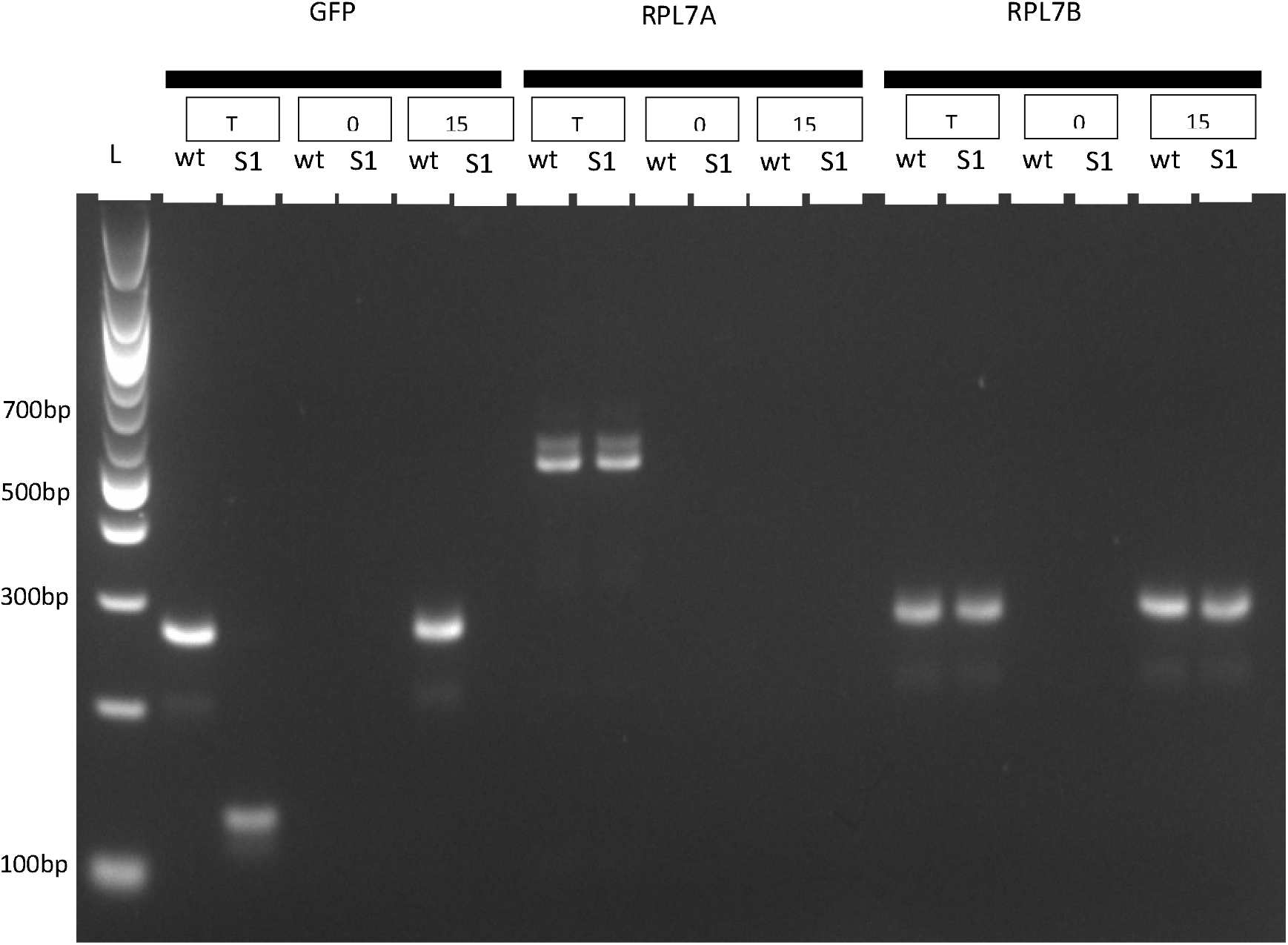
Rpl7zz protein binding. Lane 1 is a size ladder with bands of 100 bp. GFP, *RPL7A* and *RPL7B* are the three genes assayed using gene-specific primers for PCR (Supplemental Table S1). The two strains are GFP with the wild-type *RPL7B* intron 1 sequence (wt) or with the Hooks structure deletion (S1). T shows total RNA, demonstrating that the primers specifically amplify the designated gene. 0 is the RNA from the pull-down of the Rpl7-zz protein at time 0 after induction. 15 is the RNA from the pull-down of the Rpl7-zz protein at 15 minutes after induction.

## DISCUSSION

In the last several decades our knowledge of different classes of RNAs and their functional roles has been greatly expanded [43]. Some of those roles include the regulation of gene expression, and even RNAs with well-defined functions, such as snoRNAs, have additional functional roles [85, 86]. The diverse functional roles derive from the diversity of specific interactions that RNA is capable of. It can bind to RNA and DNA substrates with high specificity due to its base-pairing potential. It can also bind to proteins in a sequence-specific manner, where in addition to sequence, structure can play a significant role in the specificity of binding to proteins and other ligands, such as small molecules [30]. An important feature of RNAs is their capability of folding into two, or more, alternative structures of nearly equal free energy, and for environmental factors to influence which structure predominates. This is especially clear in the case of riboswitches, where small molecules can shift the equilibrium between two alternative structures that have altered fates for the adjacent mRNA [34, 35, 37, 87]. Riboswitches can even be sensitive to temperature to regulate gene expression [88, 89].

In the case of *RPL7B*, the predominance of the two alternative structures (Figure 4a) is determined by the concentration of Rpl7 protein. When its concentration is low, the enhancer structure predominates and leads to gene expression. When its concentration increases, the alternative structure becomes preferred and expression decreases. The need for the structural enhancer is due to the length of the intron as well as the suboptimal BP sequence. When the intron is shortened considerably, the enhancer is less important, as shown by the high expression of the U1-S1 intron variant. In every case, replacing the wild-type BP with a consensus BP increases expression and generally eliminates the need for the enhancer structure. Regulation is largely dependent on the Hooks structure, although not entirely. Even when the entire Hooks structure is deleted, there remains about a two-fold change in expression upon induction of the Rpl7 protein. The presence of the consensus BP greatly reduces the degree of repression, indicating that binding of the Rpl7 protein to the RNA does not act by blocking spliceosome access to the BP sequence. If regulation occurred by the bound protein blocking access to the BP, the wild-type and the consensus BP sequences would be equally repressed. Rather, the autoregulatory mechanism requires that the long intron has a suboptimal BP sequence so that it is inherently spliced inefficiently. The structural enhancer greatly increases its splicing efficiency so that high level expression is possible. But repression can occur when the protein is in excess, not by blocking splicing directly, but by inhibiting the formation of the structural enhancer. To our knowledge, this is a novel mechanism for autoregulation.

*RPL7A* is not regulated (Figure 1) nor bound by the Rpl7 protein (Figure 5) under the conditions of our experiments. The fact that it contains the conserved Hooks structure [73] suggests that it may be bound by the protein under alternative conditions. The *RPL7A* intron 1 proposed enhancer, or zipper stem, structure is longer and more stable [68] than our proposed structure for *RPL7B* (Figure 4), consistent with it not being bound under conditions where the *RPL7B* intron is bound. The alternative conditions under which *RPL7A* intron 1 is bound may require additional factors that help destabilize the enhancer structure.

Rpl7a and Rpl7b proteins are not functionally equivalent. They localize differently within the cell [79, 83]. *RPL7A* is the more highly expressed gene of the pair, accounting for 75% to over 90% of the Rpl7 protein [81, 90]. A strain deleted for *RPL7A* shows a significant growth defect, but deletion of *RPL7B* has no effect [81, 82]. In one study, adding a second copy of *RPL7B* did not suppress the growth defect of the *RPL7A* deletion [81], while another study reported that if the second copy of *RPL7B* is under control of the *RPL7A* promoter, that can suppress the growth defect [82]. They also respond differently to drugs. Cells expressing only *RPL7A* are more sensitive than wild-type to staurosporine, and cells expressing only *RPL7B* are less sensitive. Deletion of *RPL7B* has no effect on sensitivity to hygromycin, but cells expressing only *RPL7B* are much more sensitive. Furthermore, the two paralogs preferentially translate different subsets of genes.

Our data show that the first intron of *RPL7B* reduces gene expression even in the absence of induction of the repressor protein. When the first intron of *RPL7B* is inserted into GFP it reduces expression about 7-fold compared to the first intron (or both introns) of *RPL7A*, even in the absence of Rpl7 induction (Figure 1, uninduced curves of panels c versus g). But that reduced expression can be overcome by deleting parts of the Hooks structure required for regulation (Figure 3). Of course, Rpl7 protein, from both paralogs, exists in the cells from the normal expression of the *RPL7A* and *RPL7B* genes. That amount of protein is sufficient to reduce the expression of GFP and presumably also contributes to the reduced expression of the Rpl7b protein compared to the Rpl7a protein. These paralogs present an excellent example of ribosome specialization [91] that may explain the evolutionary advantage of maintaining both genes and their differential regulation. It will be interesting to identify conditions under which *RPL7A* is repressed and the balance of the two proteins is altered.

## Data Availability

The flow cytometry data is available on the flowrepository.org under ID FR-FCM-Z97W. Strains with the intron variants inserted into GFP are available up request.

## Acknowledgements

This work was funded in part by NIH grant R01GM140013 and partially by Washington University School of Medicine internal funds. We thank Anirudh Kesanapally for helping us decide to work on Rpl7.

