## supplemental table s1 for "Autoregulation of *RPL7B* by inhibition of a structural splicing enhancer"

**Primers used for assessing binding by Rpl7 to intron 1**

| **Primer Name** | **Sequence (5′ → 3′)** | **Target** |
| --- | --- | --- |
| RIP-GFP_cDNA | GTAAGTAGCATCACCTTCACCTTCACCGGAGACAG | GFP |
| RPL7A_cDNA | CAGCTCTTTCAGCAGCGACTTGTTCAGC | RPL7A |
| RPL7B_pull_cDNA | CTTGATTCTGACAACGAAGACCAAC | RPL7B |
| T7_hooks_structure_F | TAATACGACTCACTATAGGGTGAAGTAGTTTATCTTCTTCAAGTACC | GFP & RPL7B |
| T7_hooks_structure_R | CAGGAGTCAAGATTTTTCTATTCAAAAATG | GFP & RPL7B |
| RPL7A_Intron_F | TGGCCGCTGAGTATGTATACG | RPL7A |
| RPL7A_Intron_R | CCTTAGACTTCTTCAACTGAGATTCTGG | RPL7A |
