## supplemental figure s1 for "Autoregulation of *RPL7B* by inhibition of a structural splicing enhancer"

Fluorescence of GFP with intron 1 of *RPL7A* (top two panels) and *RPL7B* (bottom two panels) with induced proteins Rpl7a (panels 1 and 3) and Rpl7b (panels 2 and 4)

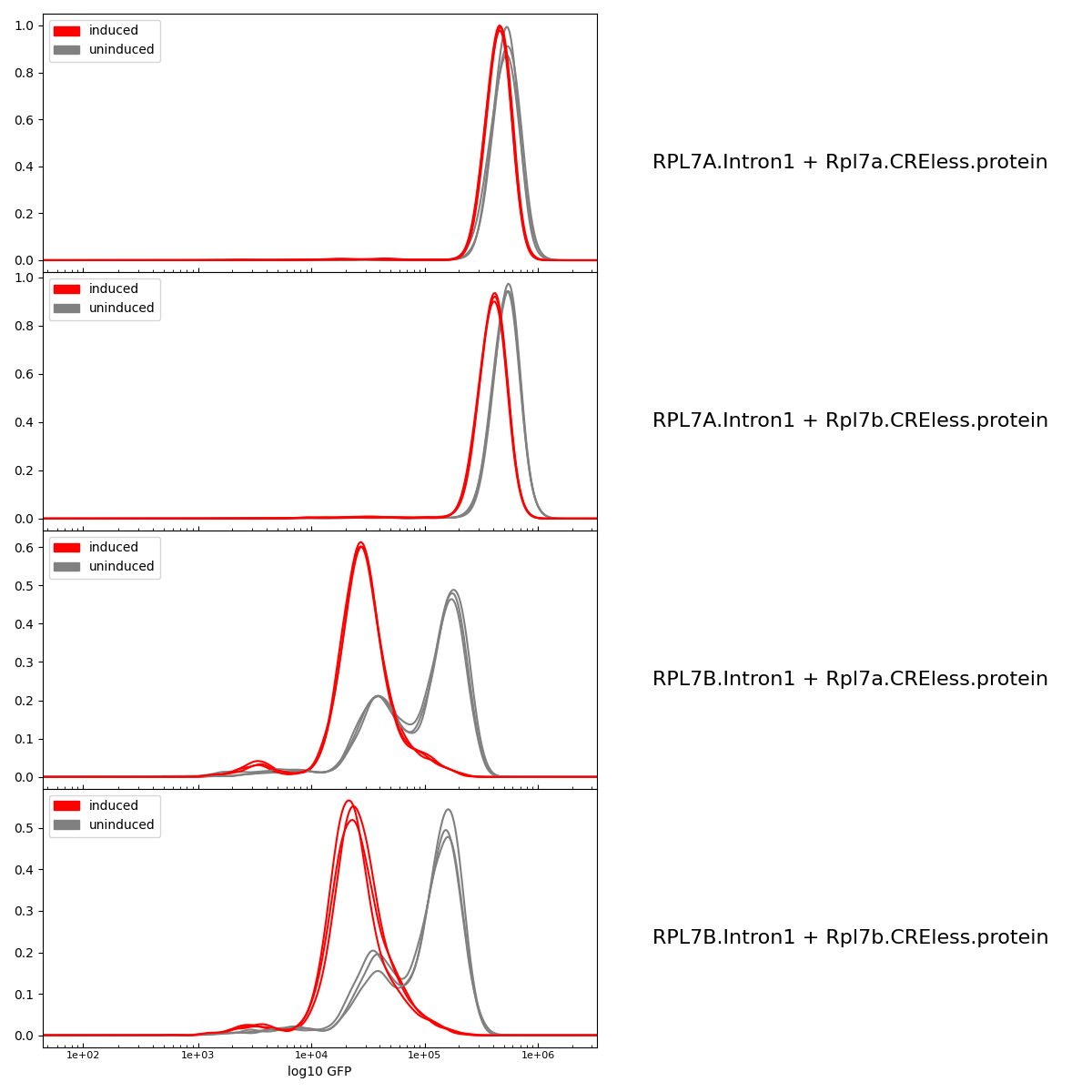
